# Factors Influencing Precision Medicine Knowledge and Attitudes

**DOI:** 10.1101/2020.06.04.133942

**Authors:** Rohini Chakravarthy, Sarah Stallings, Michael Williams, Megan Hollister, Mario Davidson, Juan Canedo, Consuelo H. Wilkins

**Author notes:** Corresponding Author Consuelo H. Wilkins, MD, MSCI, Meharry-Vanderbilt Alliance, Vanderbilt University Medical Center, 1005 Dr. D.B. Todd Jr. Blvd, Biomedical Building.

## Abstract

Precision medicine holds great promise for improving health and reducing health disparities that can be most fully realized by advancing diversity and inclusion in research participants. Without engaging underrepresented groups, precision medicine could not only fail to achieve its promise but also further exacerbate the health disparities already burdening the most vulnerable. Yet underrepresentation by people of non-European ancestry continues in precision medicine research and there are disparities across racial groups in the uptake of precision medicine applications and services. Studies have explored possible explanations for population differences in precision medicine participation, but full appreciation of the factors involved is still developing. To better inform the potential for addressing health disparities through PM, we assessed the relationship of precision medicine knowledge and trust in biomedical research with sociodemographic variables. Using a series of linear regression models applied to survey data collected in a diverse sample, we analyzed variation in both precision medicine knowledge and trust in biomedical research with socioeconomic factors as a way to understand the range of precision medicine knowledge (PMK) in a broadly representative group and its relationship to trust in research and demographic characteristics. Our results demonstrate that identifying as Black, while significantly PMK, explains only 1.5% of the PMK variance in unadjusted models and 7% of overall variance in models adjusted for meaningful covariates such as age, marital status, employment, and education. We also found a positive association between PMK and trust in biomedical research. These results indicate that race is a factor affecting PMK, even after accounting for differences in sociodemographic variables. Additional work is needed, however, to identify other factors contributing to variation in PMK as we work to increase diversity and inclusion in precision medicine applications.

## INTRODUCTION

Precision medicine (PM) is changing the one-size-fits-all healthcare paradigm for prevention and treatment of diseases through incorporation of an individual’s genetic makeup, environment, and lifestyle. (1,2) Advances in genetics, genomics, and data science have contributed to the potential for PM to reduce disease burden and mortality. At the same time, however, PM advances have the potential to widen racial and ethnic health disparities in the United States. (3–7) People of non-European ancestry are underrepresented in genetic databases, a sampling bias that can translate to clinical care bias and result in disparate outcomes or health disparities. (8,9) Little progress has been made since Need and Goldstein reviewed GWAS studies in 2009 and found that participants of European ancestry outnumbered other races 10:1. (10) As the PM initiative has expanded, we have also seen inequity in uptake of PM applications and services across racial groups.(11–13) Without significant improvement in diversity and inclusion in genomic research, any advancements from PM applications addressing the disease burden can therefore be expected to accrue inequitably, favoring those of European ancestry over those of African, Latin, and Asian ancestry. (14)

Understanding why people may be unaware of or choose not to participate in PM initiatives will be critical to the success of PM. Several explanations have been suggested for population subgroup differences in PM participation, including awareness of PM research and care options, attitudes toward PM and PM-related research topics like genetics and biobanks, trust in medical research, and socioeconomic status. Specific and accurate understanding of the factors driving the observed variation in PM knowledge and attitudes could provide evidence for how best to reduce that variation and the resulting disparities. Additionally, studies suggest that equitable PM knowledge could increase diversity among research participants. For example, in a study of research participation in a glaucoma study, African American participants with positive attitudes towards DNA donation were more likely to participate.(15) In a study of a childhood heart disease biorepository, discomfort with genetic testing was a perceived barrier to participation.(16) In a separate study, perceived benefits and positive attitudes were the most influential factors in expanding genetic cancer screening.(17) This study assesses the relationship of precision medicine knowledge (PMK) and trust in biomedical research (TBR) with sociodemographic variables using survey data collected in a large, diverse sample. The measure of PMK used includes items related to both health literacy and attitudes, factors important in the larger concept of awareness.

Awareness of PM research is significantly influenced by health literacy. Health literacy is known to increase patient participation in their own care and improve health outcomes. (18) Similarly, genomic health literacy, though less studied to date, is thought to play a role in how well a diverse and inclusive public can engage with PM, from participating in research, to processing risk assessments and acting on PM-based medical advice, to participating in policy discussions about data use and the role of PM in health care. (19) Furthermore, differences in knowledge and attitudes toward PM were influenced by health literacy more than race and ethnicity.(20,21) Researchers across disciplines have studied the contribution of the social construct of race to observed differences in PM awareness and attitudes. Familiarity with PM in general continues to be low in non-White populations in America. Awareness of the existence of genetic testing was low in African-Americans (32.9%) and Hispanics (20.6%) compared to Whites (44.4%) in a number of studies,(22–24) Realizing equity in PM knowledge and awareness should be part of a larger strategy to reduce the potential for PM to increase health disparities, as this jeopardizes its success. (25–28)

Like awareness, attitudes towards PM also differ by racial group. Minorities are more hesitant to accept PM approaches.(29) They are also less likely to support its promotion to the general public.(27) One study demonstrated that, despite similar attitudes towards the benefits of PM, Blacks were significantly more likely than Whites to be concerned about discrimination based on genetic results, use of genes and genetic information without consent, and costs to receive PM.(30)

Medical mistrust is a significant influencer of attitudes towards genetic testing.(24) In a systematic review of barriers to minority research participation, 77% of articles reviewed cited medical mistrust as a barrier which held true across all four racial minorities studied (African Americans, Pacific Islander, Latino, Asian American).(31) These differences are due to concerns that benefits will not be equitably distributed (32,33), fear of being a “guinea pig,” (34,35) and lack of legal protection for research subjects (36).

Increasingly, research suggests that factors other than race itself could explain these observed differences and therefore serve as potentially modifiable influencers of PM knowledge. Two studies reported that some recognized social determinants of health, such as income, Internet access, and numeracy skills also determined differences in PM awareness. (20,21) Preferred source of medical information may also play a key role. Previous literature suggests that Hispanics are more likely to utilize the radio for medical information, while Blacks more frequently pay attention to television.(37) Other authors have suggested that improving health information delivery to minority populations has the potential to decrease disparities in care.(38) It is possible that different racial groups have varying exposure to informative and accurate PM concepts based on their preferred sources for health information.

Using data from the Mid-South Clinical Data Research Network (MS-CDRN) Consumer Interest and Attitudes Survey, we determined the contribution of socioeconomic factors to differences in overall PM knowledge and trust in biomedical research. Other data from this survey, published elsewhere, has compared the Hall and Mainous trust scales(39) and to assessed the relationship between race, health literacy, and values important when deciding about genetic testing.(20) This current work builds upon previous literature to describe the relationship between race and overall PM knowledge. In this study, we apply validated trust scales to explain the variation in precision medicine knowledge, which we have not seen in other studies. In addition, our study creates a systematic measure of PM knowledge consisting of a calculated value for PM-related health literacy plus the six foundational PM values. This metric can be replicated in future studies to quantitatively deepen our understanding of the factors driving PM use. We hypothesized that, after controlling for age, marital status, employment, and education, race itself would not be a predictor of PM knowledge, as the social construct of race masks underlying factors contributing to disparities in PM knowledge which, unlike race, could be addressed. The findings from this work may inform efforts to recruit minority participants into PM knowledge initiatives, therefore, benefiting individual and public health.

## MATERIALS AND METHODS

### Study Population

This study was a cross-sectional survey of 3847 adult participants of the Mid-South Clinical Data Research Network (MS-CDRN), one of 11 CDRNs funded by the Patient-Centered Outcomes Research Institute. (40) The CDRN was established to facilitate involvement of patients in generating research questions and participating in research studies in order to further patient-centered research. In the MS-CDRN, clinical data from over 20 million patients are included from three large health systems encompassing 32 hospitals and hundreds of ambulatory practices. The health settings include academic medical centers, community-based hospitals, traditional outpatient clinics and federally-qualified health centers. Previous studies have included surveys of parent willingness to participate in HPV vaccination clinical trials and the relationship between depression and perceived health competence.(40) Our patient survey participants were recruited from June 2014 to June 2015 from this larger cohort. Informed consent was collected electronically prior to survey completion. The research team identified priority populations consisting of racial/ethnic minorities, individuals with multiple chronic conditions, low-income groups, rural and urban residents, and older adults. Anyone over the age of 18 with the capacity to consent was invited to participate. Recruitment strategies included in-person engagement at community health centers, minority-owned barbershops, and community health fairs. In addition, online recruitment was utilized specifically through the Vanderbilt patient portal and ResearchMatch, a volunteer registry. Participants who completed the survey received a $10 compensation. This study was approved by the Vanderbilt Medical Center Institutional Review Board.

### Survey

Patients were asked to respond to a series of demographic questions, including educational level, household income, and race. To create racial groups of sufficient size for statistical comparison, participants were asked to select one of nine racial categories. The racial groups included in this analysis were Asian, Black, Hispanic, and white.

The survey was developed based on literature evidence of common concerns about PM. The first part asked participants to rate their familiarity with PM terms on a five-point Likert scale (1=not at all familiar; 5=extremely familiar). The second part of the survey asked patients to rate the importance of certain factors in guiding future research and healthcare on a five-point Likert scale (1=not at all important; 5=extremely important). As no validated tools existed to measure these concepts at the time of this study, we developed new instruments based on previous research conducted by the study team and a literature review of genetics literacy.(41) Experts were asked to assess the relevance and clarity of each item to measure its content and face validity. The third part of the survey was one of two different validated scales to measure trust in biomedical research. Participants completed all 12 questions on the respective assigned validated scale. Half of the participants selected at random received the Hall scale,(42) while the other half received the Mainous scale.(43) This strategy was chosen to assess the consistency between the two surveys, and the results of this work are published elsewhere. (39)

In addition, basic demographic information was also collected, including ethnicity, educational attainment, and household income. Barriers to participation in research were collected using a modified, 14-item scale developed by Mouton et al.(44) For the barriers section of the survey, items “Prefer study headed by Black scientist” and “Prefer study headed by Latino scientist” were excluded due to previously observed participant discomfort with these two items. Eleven of the remaining 12 items were measured using a 5-point Likert scale based on agreement. The final item for measuring barriers to participation (“In my opinion, research in the United States is…”) had three possible response options: “ethical,” “not ethical,” and “I don’t know.” To preserve how this item loaded onto the overall score of the scale, the variable was scored such that “ethical” was recoded to 2.54, “not ethical” was recoded as 0.83, and “I don’t know” was recoded as 1.69. The total score across these 13 domains was calculated as a simple sum, with some items reversed scored so that higher scores represented a lower perception of participation barrier. The complete survey is provided in the additional supplemental material [Online Resources 1-4].

### Variables

Precision medicine knowledge (PMK) is an outcome variable created as a sum score of 10 questions. Four questions asked participants about their familiarity with PM vocabulary. The terms included genetic testing, biological indicators/biomarkers, precision medicine, and pharmacogenetics. Participants were asked to state their familiarity with the terms on a Likert Scale (1 = not at all familiar, 5 =extremely familiar). An additional six questions asked participants about attitudes towards PM concepts. Examples of questions included: “My healthcare is specific to me. No two cases are the same.”; “My genes can be used to determine the best treatment for me.”; “My genes and other health information can be used to help prevent or treat health conditions in my family,”; “My health information is kept private and secure,”; “I have access to my own health records and can decide which health care providers and researchers have access to them.”; and “I can add information about my health to my health records.” Participants were asked to rate importance on a Likert scale (1 = not at all important, 5 = extremely important). A higher total PMK score indicated greater familiarity with terms and benefits, with a maximum score of 50.

To create an overall trust in biomedical research (TBR) variable, each participant’s score from the Hall survey (42) was standardized. Items 2, 4, 7, and 10 were reverse-coded prior to summation. This was repeated for the Mainous (43) participants, with items 1-6 and 12 reverse coded prior to summation. Higher scores demonstrate higher trust towards biomedical research, with a maximum score of 5. TBR scores were treated as continuous outcome variables in the models.

### Statistical Analysis

Descriptive statistics were used to calculate demographic characteristics. For each outcome (PMK and TBR), we created a linear regression model to compare the racial group of interest and the contribution of race to the outcome. Predictive mean matching was used to account for missing data. Covariates (marital status, employment status, education, age, and gender) were selected *a posteriori*, based upon a review of relevant literature. Due to the association of race with health literacy and education, these were assessed as additional interaction variables in the model, with education coded as a continuous variable. Analysis was conducted in R (version 3.4.3).

## RESULTS

Of the 3847 participants, 83% identified as White and 15% identified as Black. The mean age was 48 (standard deviation: ±16) years, 69% percent of the population identified as female, and 61% of participants had a college degree. Table 1 contains additional demographic information.

**Table 1:**
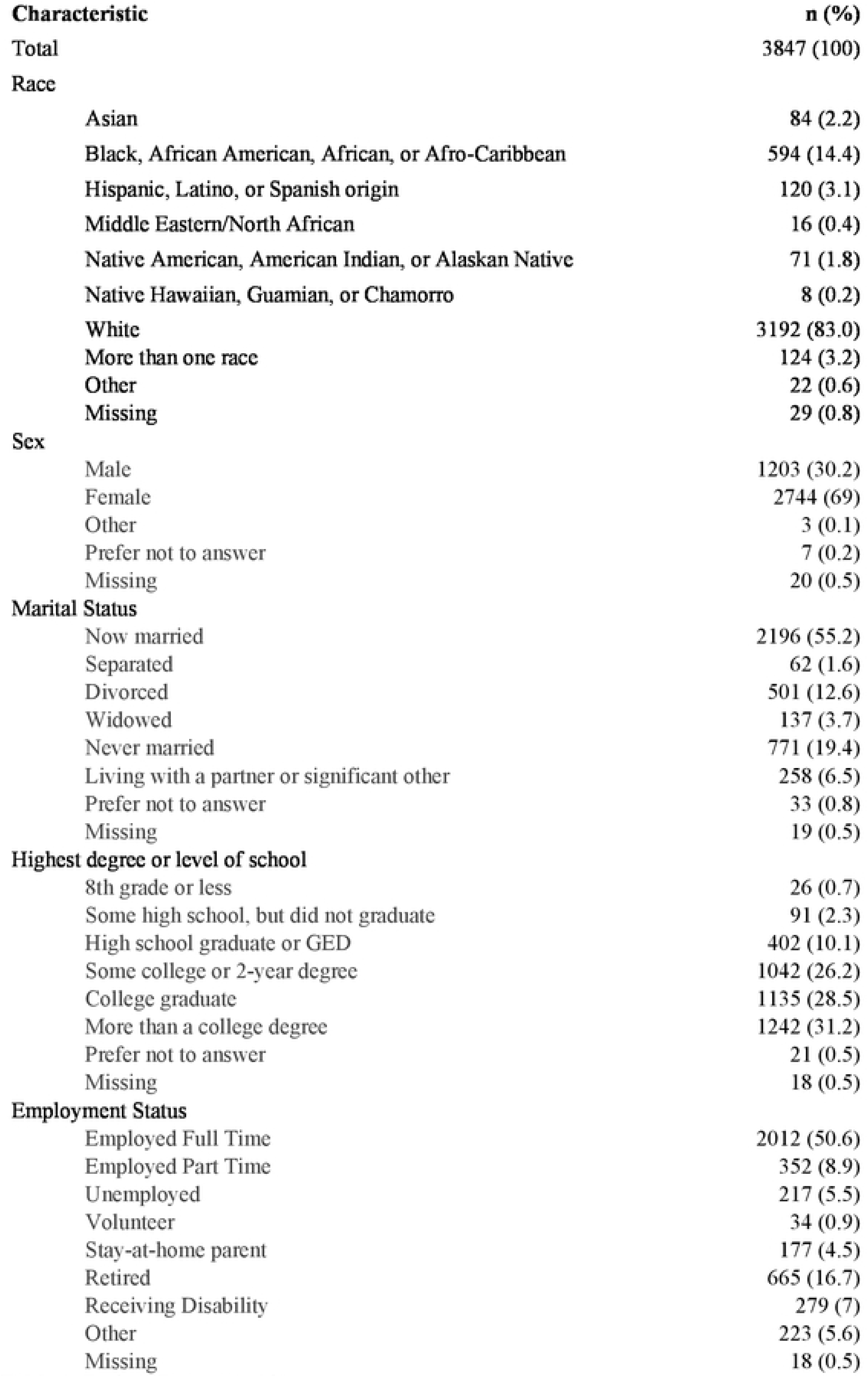
Patient Demographics

| **Characteristic** | **n (%)** |
| --- | --- |
| Total | 3847 (100) |
| **Race** |  |
| Asian | 84 (2.2) |
| Black, African American, African, or Afro-Caribbean | 594 (14.4) |
| Hispanic, Latino, or Spanish origin | 120 (3.1) |
| Middle Eastern/North African | 16 (0.4) |
| Native American, American Indian, or Alaskan Native | 71 (1.8) |
| Native Hawaiian, Guamanian, or Chamorro | 8 (0.2) |
| White | 3192 (83.0) |
| More than one race | 124 (3.2) |
| Other | 22 (0.6) |
| Missing | 29 (0.8) |
| **Sex** |  |
| Male | 1203 (30.2) |
| Female | 2744 (69) |
| Other | 3 (0.1) |
| Prefer not to answer | 7 (0.2) |
| Missing | 20 (0.5) |
| **Marital Status** |  |
| Now married | 2196 (55.2) |
| Separated | 62 (1.6) |
| Divorced | 501 (12.6) |
| Widowed | 137 (3.7) |
| Never married | 771 (19.4) |
| Living with a partner or significant other | 258 (6.5) |
| Prefer not to answer | 33 (0.8) |
| Missing | 19 (0.5) |
| **Highest degree or level of school** |  |
| 8th grade or less | 26 (0.7) |
| Some high school, but did not graduate | 91 (2.3) |
| High school graduate or GED | 402 (10.1) |
| Some college or 2-year degree | 1042 (26.2) |
| College graduate | 1135 (28.5) |
| More than a college degree | 1242 (31.2) |
| Prefer not to answer | 21 (0.5) |
| Missing | 18 (0.5) |
| **Employment Status** |  |
| Employed Full Time | 2012 (50.6) |
| Employed Part Time | 352 (8.9) |
| Unemployed | 217 (5.5) |
| Volunteer | 34 (0.9) |
| Stay-at-home parent | 177 (4.5) |
| Retired | 665 (16.7) |
| Receiving Disability | 279 (7) |
| Other | 223 (5.6) |
| Missing | 18 (0.5) |

**Table 2:** Variables: Survey questions, items, and response modes

| Variable | Questions Asked | Response |
| --- | --- | --- |
| Demographics | Age | Year of birth |
|  | Race/Ethnicity | 7 Choices + Other |
|  | Sex | Male/Female/Other |
|  | Marital Status | 5 Choices |
|  | Employment Status | 6 Choices + Other |
|  | Household Size | Free-text |
|  | Number of comorbidities | 6 Choices + Other and None |
|  | Household Income | 7 Choices |
|  | Health Insurance | 6 Choices + Other |
| Precision Medicine Vocabulary | How familiar are you with the following words or phrases?<br>Genetic testing<br>Biological indicators/biomarkers<br>Precision medicine<br>Pharmacogenetics | 5-point Likert Scale |
| Precision Medicine Attitudes | My healthcare is specific to me. No two cases are the same. | 5-point Likert Scale |
|  | My genes can be used to determine the best treatment for me. | 5-point Likert Scale |
|  | My genes and other health information can be used to help prevent or treat health conditions in my family. | 5-point Likert Scale |
|  | My health information is kept private and secure. | 5-point Likert Scale |
|  | I have access to my own health records and can decide which health care providers and researchers have access to them. | 5-point Likert Scale |
|  | I can add information about my health to my records. | 5-point Likert Scale |
| Sources of Medical Information | How much do you trust information about health or medical topics from each of the following? | 9 Choices, each rated on a 5-point Likert Scale |

**Table 3:**
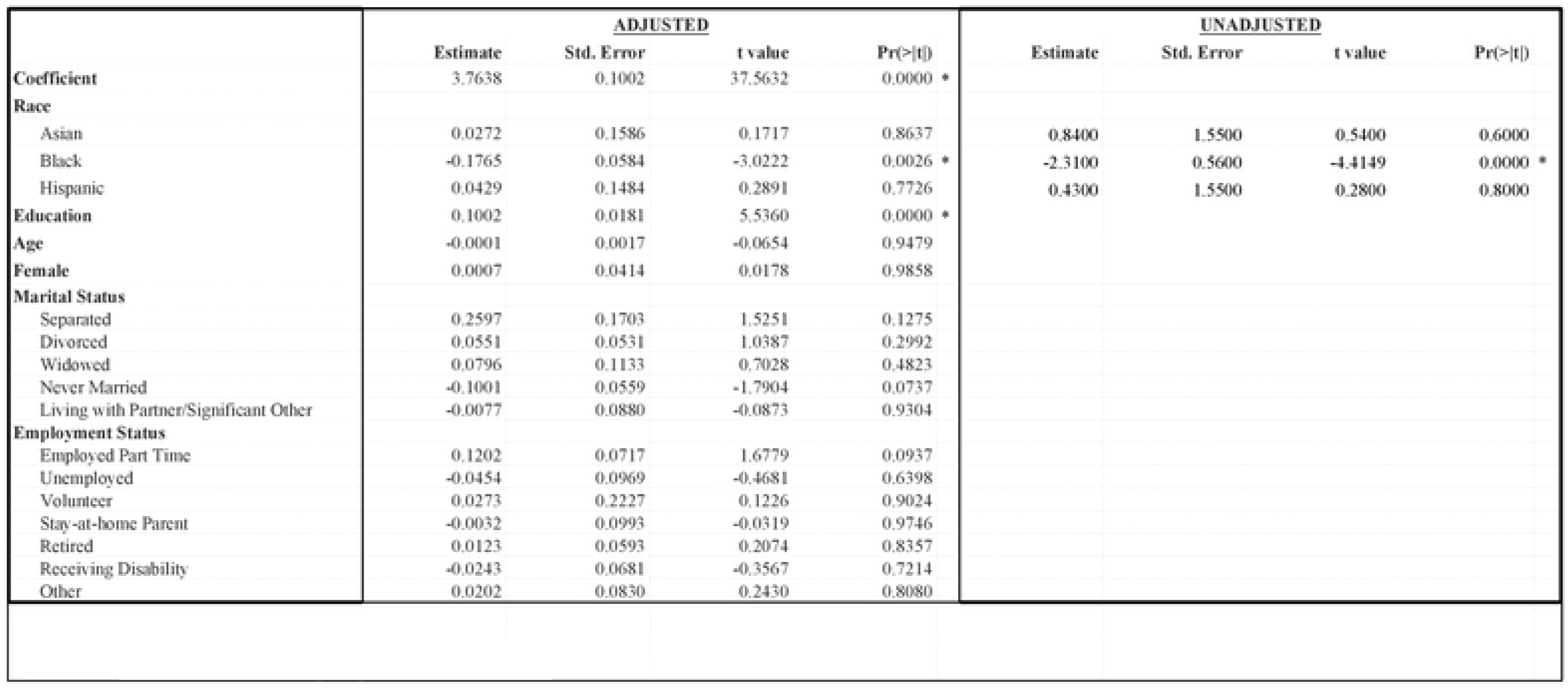
Precision Medicine knowledge Regression Model Results

### Precision Medicine Knowledge (PMK)

The average PMK score was 38.20 out of a total of 50 for the 1142 participants who fully answered the questions. Average PMK by race was 36.16 (Black), 38.36 (White), 38.87 (Asian), and 38.31 (Hispanic). Self-identification as Black was negatively correlated with PMK when compared to all other participants. This relationship describes about 1.5% of the total variation in PMK (p < 0.001). Self-identification as Asian or Hispanic was not strongly correlated with PMK. The remaining three racial categories (Middle Eastern, Native American, and Native Hawaiian) could not be compared due to small sample size. When adjusting for covariates such as age, marital status, employment, and education, self-identification as Black was still negatively correlated with PMK, with the model explaining 6.7% of overall variance (p = 0.0026). There was no evidence that race modified the effect of education on PMK when including race and education as interaction terms in the model (p = 0.2681).

### Trust in Biomedical Research (TBR)

Based on previous research using this database comparing results between the Hall and Mainous scales,(39) we were able to compare overall TBR scores for participants, regardless of which survey they took. We standardized each score distribution by subtracting its mean and dividing by its standard deviation. Self-identification as Black was negatively correlated with TBR when compared to White participants (p < 0.001). This relationship describes about 5.5% of the total variation in TBR. Self-identification as Hispanic or Asian was also negatively correlated with TBR, each explaining 0.1% of the variation. When adjusting for covariates such as age, marital status, employment, and education, self-identification as White was still positively correlated with TBR, and explained 6.8% of the overall variance.

Given that the analyzed covariates did not explain much of the variance in trust or PMK, we theorized that there may be a relationship between the two outcome variables and found a positive association between PMK and TBR (β = 0.1516, SE = 0.0512, p = 0.003).

### Preferred Health Information Source

As our initially analyzed covariates did not explain the variance in trust or PMK, we explored the theory that information source preferences could differ widely within our population and could contribute to disparities in PMK that were not addressed by the previous models. The frequency results are displayed in Table 5. When asked about preferences for health information sources, the most popular answer was internet, with approximately 32% of respondents selecting this choice. Other popular sources included doctors (31%), family (15%), or friends (10%). Among racial subgroups, the rank order for information source preference remained the same. Of the participants who answered the question and identified as White, the most frequently preferred source of information was family (30%), followed by the internet (27%), and then doctors (26%).

**Table 4:**
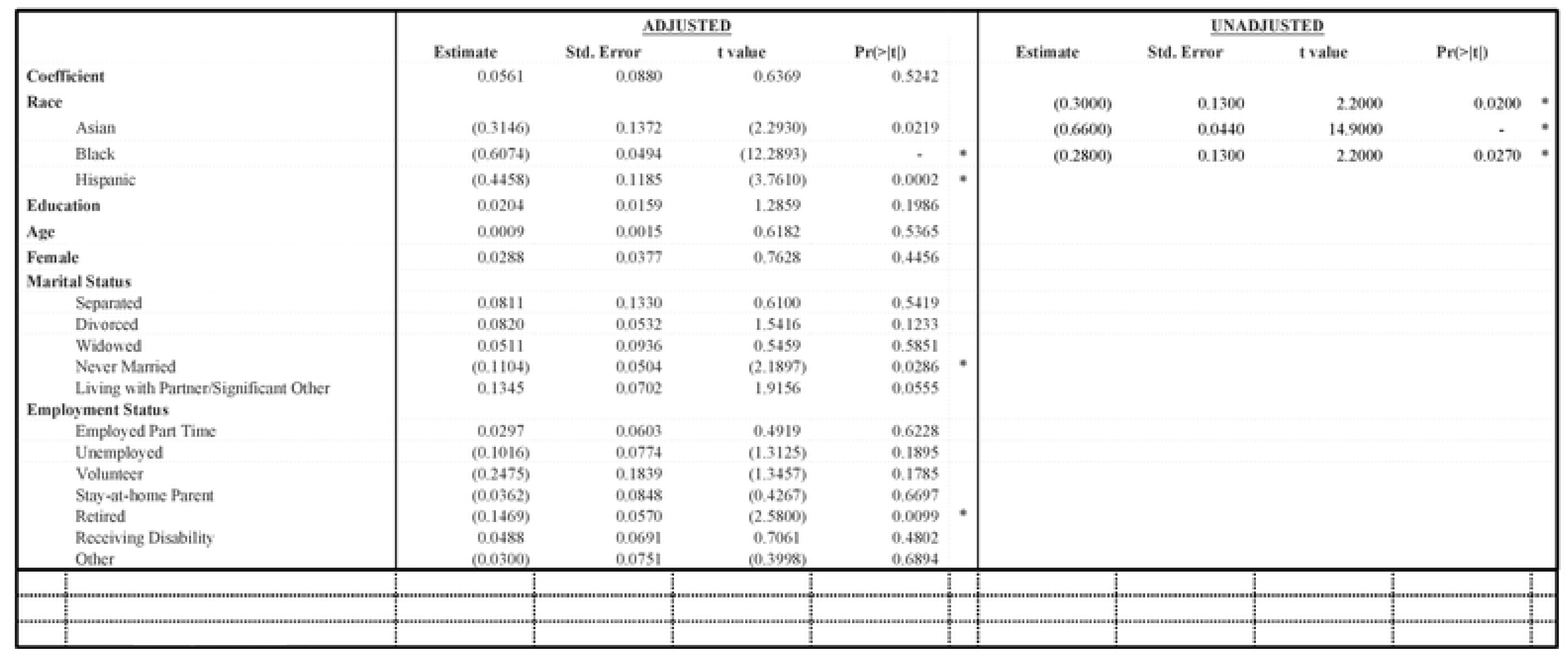
Trust Regression Models

**Table 5:** Information Source by Race

|  | Family | Friend | Doctor | Internet | Radio,<br>Newspaper,<br>Magazines | Telephone | Alternative<br>Provider | Other | Total* |
| --- | --- | --- | --- | --- | --- | --- | --- | --- | --- |
| Asian | 29% | 10% | 22% | 27% | 5% | 2% | 3% | 2% | 292 |
| Black, African American,<br>African, or Afro-Caribbean | 34% | 9% | 23% | 23% | 6% | 2% | 2% | 2% | 1765 |
| Hispanic, Latino, or Spanish<br>origin | 32% | 7% | 23% | 26% | 4% | 2% | 3% | 3% | 379 |
| Middle Eastern/North African | 25% | 14% | 22% | 22% | 5% | 3% | 5% | 6% | 65 |
| Native American, American<br>Indian, or Alaskan Native | 28% | 7% | 24% | 26% | 4% | 3% | 5% | 3% | 254 |
| Native Hawaiian, Guamanian,<br>or Chamorro | 32% | 8% | 28% | 24% | 0% | 4% | 4% | 0% | 25 |
| White | 30% | 8% | 26% | 27% | 4% | 1% | 3% | 1% | 10754 |
| Other | 40% | 4% | 20% | 29% | 2% | 0% | 2% | 4% | 55 |

\*Respondents can select more than one information source

## DISCUSSION

Identifying strategies to address disparities in PM research and applications is imperative to the successful adoption of PM strategies. Results from the MS-CDRN Participant Survey, designed to collect data on stakeholder opinions about participation in that research network, have allowed us to identify gaps between race, socioeconomic barriers, and perceived mistrust on overall PMK. Black participants have significantly lower PMK and TBR, even after controlling for meaningful covariates. This is also true of Asian and Hispanic participants for TBR models. However, as the models explain less than 7% in the overall variation in PMK, even after adjusting for meaningful covariates included in this survey, indicating that the survey data are an incomplete measure of factors related to PMK and TBR. Moreover, there is a significant association between PMK and TBR. The magnitude of these differences appears to be greater for Black participants compared to Hispanic or Asian participants. Additionally, the most popular health information sources (family, internet, and doctors) were consistent, regardless of race, and therefore are unlikely to contribute to the differences observed.

These results refuted our original hypothesis that the effects of race would be null after accounting for sociodemographic variables of education, income, employment, and marital status and confirms previous research citing race as a barrier for participation in biomedical research. The results from the models created in this study suggest that we still have an incomplete picture of factors influencing PMK. Until we have an understanding of why participants choose to not participate in PM initiatives or have not yet accepted the potential benefits, disparities in PM acceptance will grow. These factors could include previous interactions with the healthcare system,(43) self-efficacy,(45) and health literacy, specifically genetics literacy.(46) While our survey captures many social determinants of health, it is possible that additional drivers such as food insecurity, housing instability, or neighborhood safety issues may limit the benefits of PM in certain populations.

PM participation is modifiable, and early adopters have consistent personality traits, regardless of minority status.(47) Meaningful interventions, such as better representation of socioeconomic diversity in research leadership, tailored health education materials of appropriate literacy, and improved genetics education of the public, are all potential methods to decrease disparities in PM knowledge and attitudes which, in turn, could decrease differences in participation.(14) Furthermore, interventions that address logistical barriers, such as the complexity of payer coverage for genetic testing,(48) the value of negative or unknown results to patients,(49,50) or the burden on primary care providers,(51) is sorely needed to increase potential benefits of PM.

Previous research also supports these findings. A qualitative study found that African American focus group participants recommended a reduction in technical detail for genetics communication aids.(52) Both providers(53) and patients(54) recognize the need for increased genetics education to alleviate disparities in knowledge and attitudes towards PM. Culturally-tailored material and engagement of local stakeholders has been successful in improving recruitment of underrepresented groups in research.(55) Among Asian Americans, opportunities for improving participation include providing linguistically appropriate materials on genetic testing.(56) In addition, more granular details including specific countries of origin and length of time in the US could help researchers better understand differences within the Asian population as a whole.(57)

In response to these and other studies, there is significant ongoing work to specifically address disparities in PM. One such example is the All of Us Research Program, which aims to increase genetic biodiversity by creating a national database.(58) This organization offers materials in a variety of different languages, has partnered with trustworthy community organizations to engage and retain participants, utilized a patient advisory board early to advise development.(59) Meanwhile, The Personalized Medicine Research Project consulted patient participants in the design of the protocol to include changes such as newsletters to disseminate information to study participants, external advisory boards, and focus groups.(60) We believe that engaging participants in these novel ways can mitigate differences in PMK and TBR though the results of these initiatives are still being studied.

A strength of the current study is that the demographic distribution is similar to the MS-CDRN population as a whole. Special efforts were made to recruit underrepresented groups to participate in this MS- CDRN Survey through recruitment at federally-qualified health centers and community settings (i.e. barber shops). The population’s median age was reported to be 56.3. 84.1% were White, 9.9% were Black, 1.8% were Hispanic, and Asian was not reported in the results. A predominance of females was also seen in the study population as a whole (64%).(61) The surveys were comprehensive, including information on several variables impacting participation in PM, such as attitude, awareness, and trust in medical research, as well as several other confounding variables including age, marital status, employment, and education, which were successfully incorporated into statistical models. Increasing the length of the survey to explore additional topics such as use of genetic testing by law enforcement or access to direct-to-consumer testing were not directly measured by the validated tools used in this study.

### Limitations

There are several limitations to this study. First, it relied on self-reported data and not objective measurements of PM knowledge. As no validated tools exist, the authors relied on methods that have been utilized in other genetics literature. Second, the sample was limited to those who electively participated in a survey about research attitudes. This resulted in a participant cohort with above average levels of education and income that was majority female. In addition, this nonresponse bias may have led to higher participation rates among people with higher TBR scores compared to the general population. For these reasons, these conclusions may not be generalizable to the whole population. Unfortunately, this study was not powered to address the nuances within race and can only draw conclusions generally about those who identify broadly as White, African American, Asian, and Hispanic. These categories do not fully represent the spectrum of racial diversity.

## CONCLUSIONS

As the field of PM grows, we must be cautious about growing disparities in its understanding and dissemination. While statistical models are able to capture individual factors such as education, income, and race it is clear that PM knowledge represents the end result of a complex, longitudinal interaction between providers, patients, the scientific community, society, and the environment. Future statistical models must incorporate additional factors to better understand variations in PM knowledge/awareness. Efforts must be made to address sociocultural barriers as well as logistical barriers to accessing genomic testing. Equitable distribution of PM interventions and their benefits requires deepening our genomic knowledge in addition to our sociocultural knowledge.

## ACKNOWLEDGEMENTS

The authors would like to thank the researchers, recruiters, and survey participants throughout the mid-south that allowed us to conduct this research. Thanks to Dr. Victoria Villalta-Gil and Donna Ingles for their critical review of this manuscript.

